# Robust and sensitive GFP-based cGMP sensor for real time imaging in intact *Caenorhabditis elegans*

**DOI:** 10.1101/433425

**Authors:** Sarah Woldemariam, Jatin Nagpal, Joy Li, Martin W. Schneider, Raakhee Shankar, Mary Futey, Aruna Varshney, Kristine Andersen, Benjamin Barsi-Rhyne, Alan Tran, Wagner Steuer Costa, Chantal Brueggemann, O. Scott Hamilton, Denise M. Ferkey, Miri VanHoven, Alexander Gottschalk, Noelle L’Etoile

## Abstract

cGMP is a ubiquitous second messenger that plays a role in sensory signaling and plasticity through its regulation of ion channels and kinases. Previous studies that primarily used genetic and biochemical tools suggest that cGMP is spatiotemporally regulated in multiple sensory modalities, including light, heat, gases, salt and odor. FRET- and GFP-based cGMP sensors were developed to visualize cGMP in primary cell culture and *Caenorhabditis elegans* to corroborate these findings. While a FRET-based sensor has been used in an intact animal to visualize cGMP, the requirement of a multiple emission system limits its ability to be used on its own as well as with other sensors and fluorescent markers. Here, we demonstrate that WincG2, a codon-optimized version of the cpEGFP-based cGMP sensor FlincG3, can be used in *C. elegans* to visualize rapidly changing cGMP levels in living, behaving animals using a single fluorophore. We coexpressed the sensor with the blue light-activated guanylyl cyclases BeCyclOp and bPGC in body wall muscles and found that the rate of WincG2 fluorescence correlated with the rate of cGMP production by each cyclase. Furthermore, we show that WincG2 responds linearly upon NaCl concentration changes and SDS presentation in the cell bodies of the gustatory neuron ASER and the nociceptive phasmid neuron PHB, respectively. Intriguingly, WincG2 fluorescence in the ASER cell body decreased in response to a NaCl concentration downstep and either stopped decreasing or increased in response to a NaCl concentration upstep, which is opposite in sign to previously published calcium recordings. These results illustrate that WincG2 can be used to report rapidly changing cGMP levels in an intact animal and that the reporter can potentially reveal unexpected spatiotemporal landscapes of cGMP in response to stimuli.

**Author Summary:** cGMP is a second messenger that plays an important role in sensory signaling and neural plasticity. Previous genetic and biochemical studies indirectly suggest that cGMP is spatiotemporally regulated in neurons to modulate neural activity. While a FRET-based sensor for cGMP has been used in intact *Caenorhabditis elegans* to examine its spatiotemporal regulation in neurobiological processes, its use has been limited due to the complicated setup required to image this type of sensor. Here, we describe a GFP-based cGMP sensor that has been codon optimized for use in *C. elegans* and demonstrate that it responds robustly and reliably to endogenously changing cGMP levels. We show that the sensor responds to cGMP production by coexpressing it with blue light-activated guanylyl cyclases, and we show that it responds to NaCl and sodium dodecyl sulfate when expressed in a gustatory and nociceptive neuron, respectively. We think that this sensor can be used to investigate the spatiotemporal regulation of cGMP in neurons and its relationship to neural activity.

## Introduction

The canonical second messenger molecule cGMP (cyclic guanosine monophosphate) regulates richly diverse functions in an animal’s nervous system. cGMP signaling underlies the outgrowth of axons and the transduction of light, scent and other environmental cues to electrical signals in the brain [1]. Because so many neurobiological processes revolve around cGMP, having a robust, easy to use visual reporter for cGMP with precise temporal and spatial fidelity is critical to complement the primarily pharmacological, biochemical and genetic approaches used to study this second messenger’s role in these processes. Such a reporter can be used to illuminate how producers and regulators of cGMP shape the landscape of this cyclic nucleotide in neurons.

Since cGMP is used in diverse cell types as a second messenger, its levels need to be regulated in ways that serve the cells’ distinct functions. cGMP production can be regulated directly by stimuli such as ions, peptides, temperature and gases that interact with guanylyl cyclases (GCs) that convert GTP to cyclic GMP [2–8]. Recent evidence suggests that stimuli such as ions, peptides, Gα and temperature appear to largely regulate transmembrane receptor guanylyl cyclases (rGCs), while membrane permeable gases such as nitric oxide, carbon dioxide and oxygen have been shown to directly activate soluble guanylyl cyclases (sGCs) [6–8]. Opposing the activity of GCs are phosphodiesterases (PDEs) that hydrolyze cGMP; they can be regulated by cGMP, cAMP, Ca^2+^, kinases and the γ subunit of heteromeric G proteins [9–11]. cGMP effectors, including cyclic nucleotide-gated channels and kinases, can act rapidly by changing the membrane potential of a cell (e.g. the visual system); they can also have slower, more long lasting effects on gene expression [12,13]. Thus, the precise subcellular localization of GC and PDE proteins and their temporally regulated activities are likely to produce a complex and dynamic landscape of varying cGMP levels and restricted localized activation of effector proteins.

In neurobiology, the spatial localization of cGMP signal transduction pathways suggest that spatiotemporal regulation of cGMP could play a role in sensory transduction and plasticity, which is likely to be different in distinct neurons [14,15]. To this end, the transparent nematode *C. elegans* is an ideal model system to visualize cGMP in neurobiological processes. While the use of genetic manipulations in this animal yielded valuable insights into the role of cGMP in sensory transduction, a visual tool that complements this approach has the potential to reveal how this neuromodulator is spatiotemporally regulated in real time. For instance, while mutants lacking distinct, functional rGCs revealed that cGMP signaling was required for sensing specific gustatory cues and NaCl concentration cultivation preference in *C. elegans*, it is unclear whether this second messenger is spatiotemporally regulated [3,4,16]. Additionally, while genetic evidence suggests that the cGMP-dependent protein kinase EGL-4 and GCs regulate *C. elegans’* sensitivity to quinine through the flow of cGMP from sensory neurons to the nociceptive neuron ASH through gap junctions [17,18], a visual tool could greatly enhance our understanding of the process and its dynamics. In another nociceptive neuron, PHB, genetic evidence suggests that the G protein coupled receptor *srb-6* is required to sense noxious liquids including sodium dodecyl sulfate (SDS) and dodecanoic acid [19], and calcium recordings suggest that avoidance of isoamyl alcohol is at least partially mediated by the cGMP-gated cation channel TAX-2/TAX-4, both suggesting that cGMP flux is required for sensing some nociceptive cues [20]. Thus, genetic tools demonstrate the importance of cGMP signaling in distinct sensory modalities, and a tool to visualize cGMP fluxes with precise temporal and spatial fidelity could deepen our understanding of these important processes, offering a more complete picture of the cGMP landscape and dynamics in cells. Such a tool will provide an essential complementary approach to the primarily genetic approaches that have been used to examine cGMP dynamics in this animal [21].

Though a Förster resonance energy transfer (FRET)-based tool has been used to this end, a single channel fluorophore tool provides additional flexibility as it would allow for more wavelengths to be used, making it more amenable for visualization with other reporters (e.g. calcium sensors and fluorescent markers for organelles). In AWC, this FRET-based sensor showed compartmentalized cGMP dynamics in response to odorants [22]. Additionally, in PQR, simultaneous imaging of the cGMP sensor with a calcium sensor in response to a 7% to 21% increase in O_2_ revealed that a decrease in cGMP correlated with an increase in Ca^2+^, suggesting potential compartmentalization of cGMP dynamics [8]. While these studies demonstrate that the FRET-based sensor can be used to visualize the cGMP landscape in neurons, the complexity of a multiple emission system might raise a barrier to its use. Thus, having a robust, single fluorophore sensor for cGMP will complement the use of calcium sensors in this animal, providing a way to investigate how the spatiotemporal regulation of cGMP influences neural activity *in vivo*. Such a tool would be maximally efficient for probing the interplay between cGMP and calcium dynamics within sensory compartments in this transparent organism.

Here, we show that WincG2 (**W**orm **In**dicator of **cG**MP – **2**), the *C. elegans* codon-optimized version of the GFP-based circularly permuted cGMP sensor FlincG3, reports cGMP dynamics *in vivo* in *C. elegans* [23]. We characterize the biochemical and biophysical properties of WincG2 upon cGMP binding *in vivo* by ectopically expressing the blue light-activated guanylyl cyclases BeCyclOp and bPGC in muscle cells [24]. Using the WincG2 reporter, we show for the first time that sensory stimulation of either gustatory or nociceptive neurons triggers cGMP changes. We demonstrate that cGMP reliably decreases in response to NaCl concentration downsteps and increases in response to NaCl concentration upsteps in the salt-sensing neuron, ASER, which is opposite in sign to calcium transients that increase in response to NaCl concentration downsteps [3,25,26]. Additionally, we demonstrate that cGMP flux can be observed in the phasmid PHB neurons in response to a repulsive stimulus. Our results demonstrate that the GFP-based WincG2 is a versatile tool for the study of cGMP dynamics in different sensory modalities in intact animals using a single fluorophore.

## Results

### WincG2 is a codon-optimized circularly permuted cGMP sensor derived from the mammalian sensor, FlincG3

WincG2, a genetically-encoded, circularly permuted EGFP-based cGMP sensor, is the *C. elegans* codon-optimized version of FlincG3, which was initially characterized both *in vitro* and in cell lines [23]. FlincG3 is based on FlincG, an earlier version of the sensor. Like FlincG, FlincG3 contains the N-terminal region of protein kinase G (PKG) I α, which is comprised of two cGMP-binding domains that bind cGMP cooperatively. The first 77 amino acids of the N-terminal region of the cGMP-binding domain were deleted to prevent interactions with endogenous PKG [27]. This region of PKG I α is appended to the N-terminus of circularly permuted EGFP (cpEGFP) [23,27]. In the presence of cGMP, FlincG3 fluorescence increases, presumably due to the conformational changes of the sensor upon cGMP binding that allow the beta barrel of GFP to form and create the appropriate environment for fluorophore maturation [23,27]. The response amplitude of the FlincG3 sensor to cGMP was enhanced by a M335K substitution located outside the beta barrel of the cpEGFP domain (Fig 1) [23]. FlincG exhibits rapid kinetics, and FlincG3 retains this property as it rapidly detects changes in endogenous cGMP levels in the nanomolar to low micromolar range in response to nitric oxide when expressed in HEK_GC/PDE5_ cells [23,27]. Additionally, FlincG3 fluorescence increases *in vitro* in response to a 230-fold lower concentration of cGMP than cAMP, suggesting that it preferentially binds to cGMP [23]. For our study we codon optimized FlincG3 for *C. elegans* and inserted it into a standard *C. elegans* expression vector (Fig 1).

**Fig 1.**
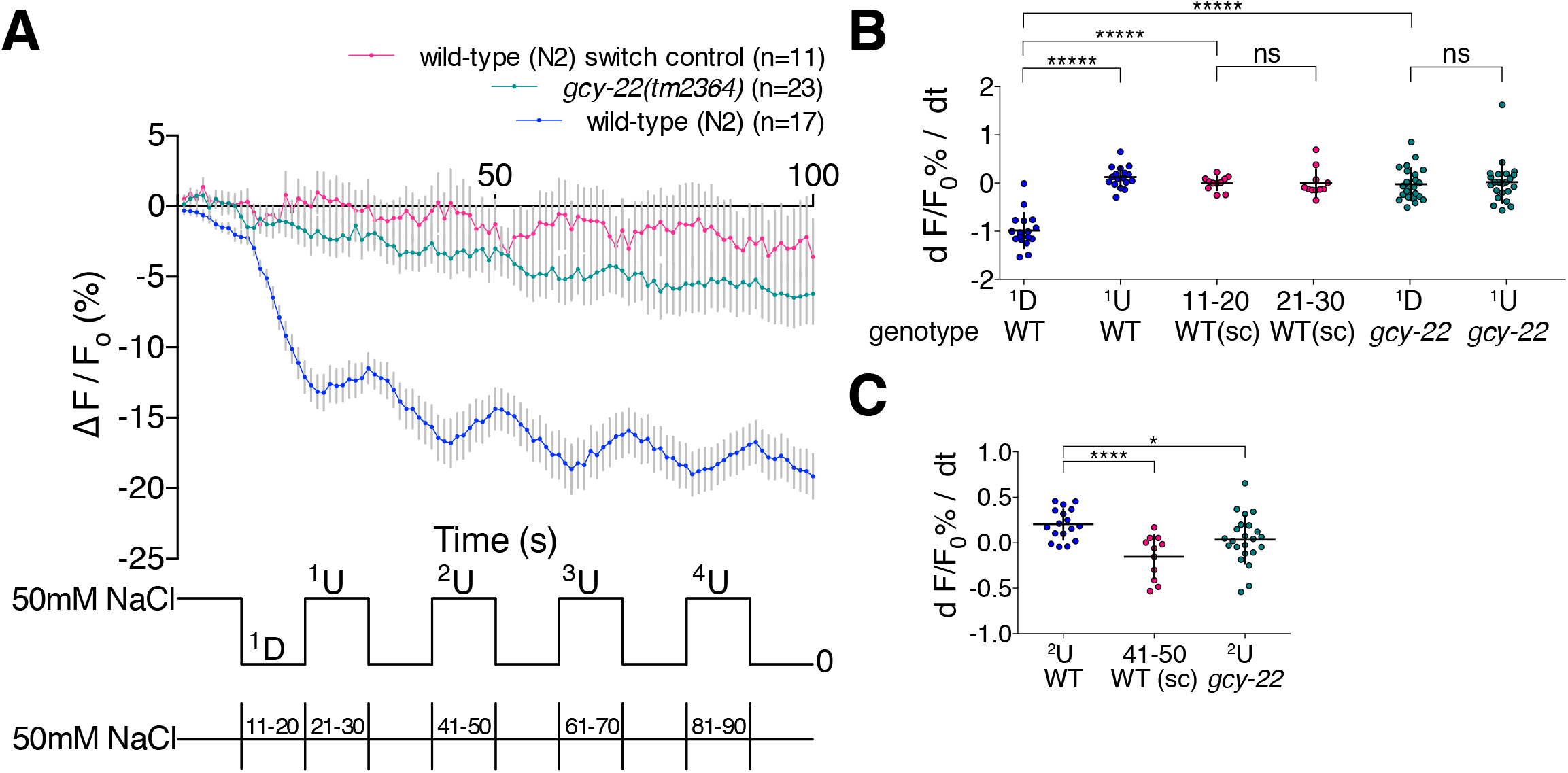
WincG2 is a GFP-based cGMP sensor codon optimized for use in *C. elegans*. WincG2 is the *C. elegans* codon-optimized version of the mammalian cGMP sensor FlincG3 (figured adapted from Garthwaite) [23]. This GFP-based sensor contains two in-tandem protein kinase G (PKG) I α cGMP binding domains that bind cGMP cooperatively (PKG1α (77–356); maroon); this regulatory PKG domain is attached to the N terminus of circularly permuted EGFP (cpEGFP; green). Changing the methionine at position 335, located outside the beta barrel of the cpEGFP domain, to lysine (M335K), improved the response amplitude of the sensor to cGMP [23]. GGTGGS is a linker between the two GFP halves. This linker, along with the 6xHis-tag region (H6) and the Tag Region, were retained from FlincG3. This *C. elegans* codon-optomized sensor, prepared by Genscript, was inserted into a worm-specific Fire vector, pPD95.75, which contains synthetic introns (SynIVS.A and SynIVS.L; blue) to facilitate expression, a multiple cloning site (MCS) and the 3’ untranslated region of *unc-54* (*unc-54* 3’ UTR; orange).

### Stimulation of blue light-activated guanylyl cyclases increases WincG2 fluorescence

To test whether WincG2 can detect rapid changes in cGMP levels in an intact animal, we utilized the *C. elegans* body wall muscle cells, which lack most endogenous GCs. We coexpressed the reporter along with heterologous light-inducible GCs that have different cGMP production rates [24,28]. BeCyclOp is a microbial rhodopsin from *Blastocladiella emersonii* that is linked to a cytosolic GC domain (Fig 2A). It detects photons by absorption using the retinal chromophore and transmits this signal into activation of the GC domain [24]. bPGC *(Beggiatoa sp*. photoactivated guanylyl cyclase) is a BLUF-domain photosensor that is coupled to a GC domain (Fig 2C). It originates from bPAC *(Beggiatoa sp*. photoactivated adenylyl cyclase) that was mutated to generate cGMP rather than cAMP [28].

**Fig 2.**
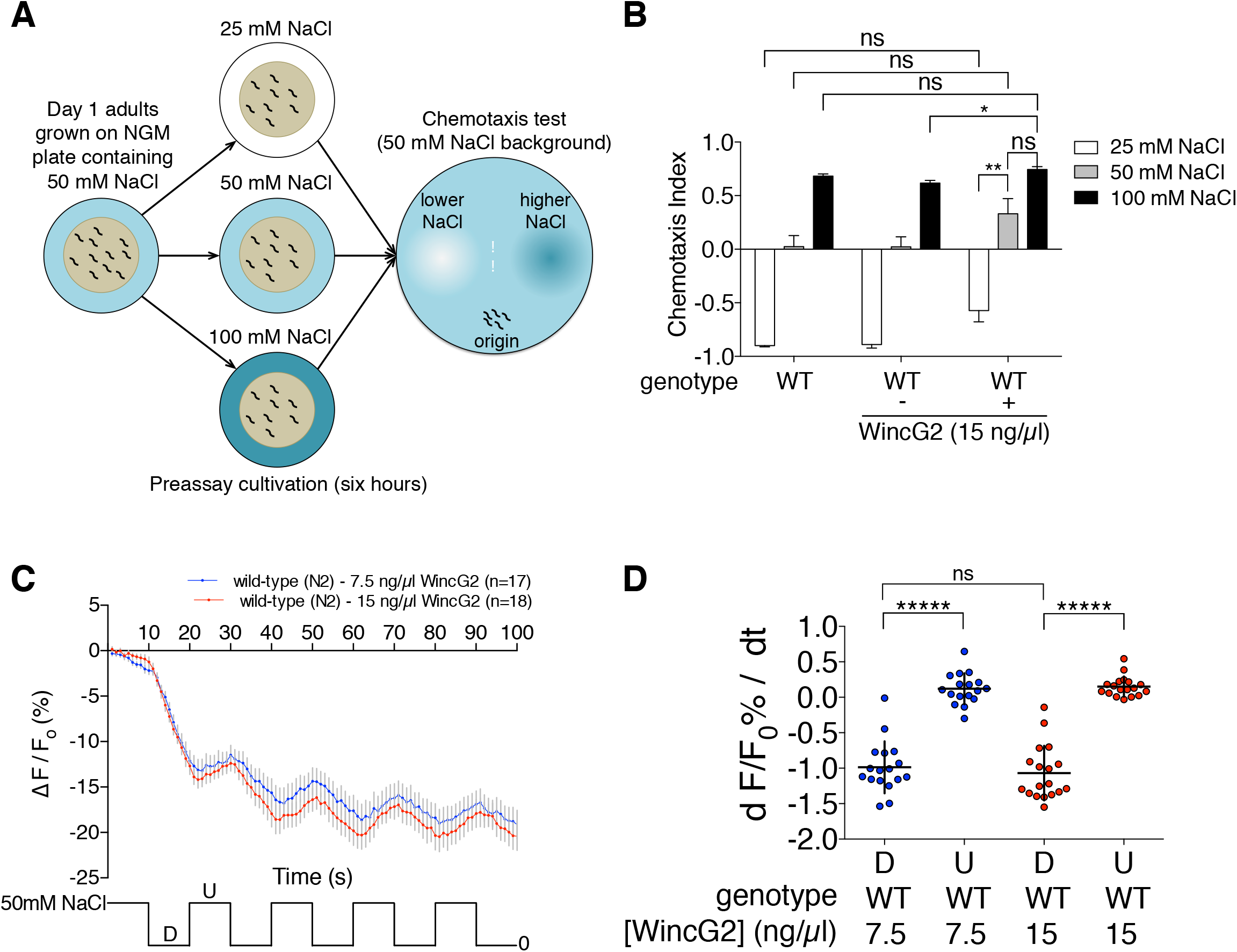
WincG2 fluorescence increases upon stimulation of blue light-activated guanylyl cyclases when coexpressed in body wall muscle cells. (A) BeCyclOp is a fungal blue light-activated guanylyl cyclase that generates cGMP with a turnover rate of approximately 17 cGMP s^−1^ at 20°C [24]. (B) (F-F_0_)/F_0_ for WincG2 fluorescence intensity in *myo-3p*::CyclOp::SL2::mCherry; *myo-3p*::WincG2 animals grown in the absence and presence of all*-trans* retinal (ATR). n=6 for WincG2 fluorescence intensity without ATR (light green, bottom); n=5 for WincG2 fluorescence intensity with ATR (dark green, top). Inset shows raw traces and indicates F_0_. (C) bPGC is a bacterial blue light-activated guanylyl cyclase containing a BLUF (blue light receptor using FAD) domain with an estimated turnover rate of 0.2 cGMP s^−1^ [28]. (D) (F-F_0_)/F_0_ for WincG2 fluorescence intensity in *myo-3p*::bPGC::SL2::mCherry; *myo-3p*::WincG2 animals (n=5, 27 ROIs). Inset shows raw traces and indicates F_0_. (E) bPAC is a bacterial blue-light activated adenylyl cyclase containing a BLUF (blue light receptor using FAD) domain. *In vitro* cAMP production in the presence of blue light is 10 ± 2 nmol cAMP per minute per mg [31]. (F) (F-F_0_)/F_0_ for WincG2 fluorescence intensity in *myo-3p*::bPAC::SL2::mCherry, *myo-3p*::WincG2 animals. n=7. Inset shows average of traces during the first two seconds of recording.

To test whether changes in WincG2 fluorescence and dynamics correspond with cGMP production by BeCyclOp, animals coexpressing WincG2 and BeCyclOp were grown with or without all*-trans* retinal (ATR), which is required for BeCyclOp activity. When WincG2 and BeCyclOp were coexpressed in body wall muscle cells in the presence of ATR, an acute increase in WincG2 fluorescence (peak F-F_0_/F_0_ = 0.218 ± 0.023 at 0.49s) was observed upon continuous blue light illumination, which activates BeCyclOp. This was followed by a slight decay over the duration of the recording (Fig 2B: top green trace). By contrast, animals grown without ATR and thus having no BeCyclOp activation exhibited an apparent decrease in WincG2 fluorescence (F-F_0_/F_0_ plateaued at approximately −0.310 to −0.312 beginning at 8.98s) when exposed to blue light (Fig 2B: bottom blue trace). The initial signal decayed; F-F_0_ became negative, then plateaued and remained steady for the duration of the recording. Notably, the initial fluorescence intensity F_0_ (as measured in the absence of the rhodopsin cofactor ATR; bottom blue trace in Fig 2B) exhibited a rapid drop, possibly due to photoswitching behavior that was previously observed for other fluorescent proteins [29,30]. Taken together, we interpret these results to indicate that WincG2 fluorescence correlates with activation of BeCyclOp by blue light.

bPGC, a blue light-activated GC derived from the corresponding adenylyl cyclase bPAC (also known as BlaC), produces 50-fold less cGMP per unit time relative to BeCyclOp [24,28]. WincG2 fluorescence increased in the order of minutes upon continuous activation of bPGC with blue light (peak F-F_0_/F_0_ = 0.122 ± 0.023 at 145.2 s) (Fig 2D). Note that at the onset of blue light illumination, WincG2 fluorescence increased, then decreased rapidly; this is presumably the same rapid photoswitching observed in the BeCyclOp experiment (the time constants for decay of the signal were essentially identical: 0.383 s for the experiment in Fig 2B, and 0.297 for the experiment in Fig 2D, in line with the hypothesis that this due to the same photophysical process). We chose F_0_ after this photoswitching at 1 second after light onset (Fig 2D: inset). At this time, meaningful amounts of cGMP begin to develop, as judged from experiments in which bPGC was coexpressed in body wall muscle with the cGMP-gated cation channel TAX-2/TAX-4; muscle contractions from ion influx begin to be observable after 1 second of blue light (S1 Fig).

After photoswitching, we observed a slow rise in WincG2 fluorescence, which we interpret to be due to the slower kinetics of bPGC relative to BeCyclOp. By contrast, WincG2 fluorescence increased acutely upon activation of BeCyclOp, suggesting that the rate of change of WincG2 fluorescence correlates with the rate of cGMP production by each GC.

Bhargava *et al*. showed that FlincG3 has a 230-fold lower EC50 for cGMP relative to cAMP [23]. To assess whether WincG2 fluorescence changes with increasing cAMP levels *in vivo*, WincG2 was coexpressed with bPAC, a bacterial blue light-activated adenylyl cyclase, in body wall muscle cells (Fig 2E) [31]. Following a fast drop in fluorescence, these animals showed a 10% increase in WincG2 fluorescence upon blue light stimulation of bPAC that peaked and decayed in a manner similar to that of the WincG2 response to BeCyclOp, albeit with slightly slower kinetics (Fig 2F). Thus, WincG2 appears to respond to cAMP. Indeed, bPAC is an efficient adenylyl cyclase that produces cAMP at the rate of 10 ± 2 nmol per minute per mg. Thus, it is not surprising that WincG2 responds to the high production of cAMP by bPAC [31]. Since there are no amino acid changes between WincG2 and FlincG3, it is expected that WincG2 is also activated more effectively by cGMP relative to cAMP. However, these results indicate that it is important to control for WincG2 response to cAMP.

### WincG2 fluorescence in ASER cell body changes in response to NaCl concentration steps and depends on the rGC GCY-22

Genetic and calcium imaging studies indirectly suggest that cGMP in the gustatory neuron ASER mediates both acute sensation of NaCl removal and the food-mediated gradual resetting of NaCl cultivation preference [16,25]. ASER expresses multiple rGCs and may use cGMP to gate the cyclic nucleotide-gated TAX-2/TAX-4 cation channel upon NaCl concentration changes [3]. Consistent with this hypothesis, NaCl downsteps trigger an influx of calcium into ASER which was blocked in animals lacking TAX-2 or TAX-4 [25]. Additionally, a study indicating that cGMP could be a putative second messenger in ASER revealed that loss of the rGC GCY-22 blunts chemotaxis to Cl^−^ [4]. This suggests that cGMP levels could be modulated by changes in NaCl concentration [3].

To explore this hypothesis, we expressed WincG2 in the ASE neuron pair and monitored the sensor’s response in the ASER cell body to ten 10-second steps between 50 and 0 mM NaCl. WincG2 fluorescence decreased linearly (R^2^ = 0.9962) in response to a 50 to 0 mM NaCl downstep and stopped decreasing in response to the first 0 to 50 mM NaCl upstep (Fig 3A: blue trace, bottom). The slopes between the first downstep and upstep are different (p<0.00001; see Materials and Methods for statistical analysis), suggesting that WincG2 responds quickly to changing NaCl concentrations (Fig 3B: first pair, blue).

**Fig 3.**
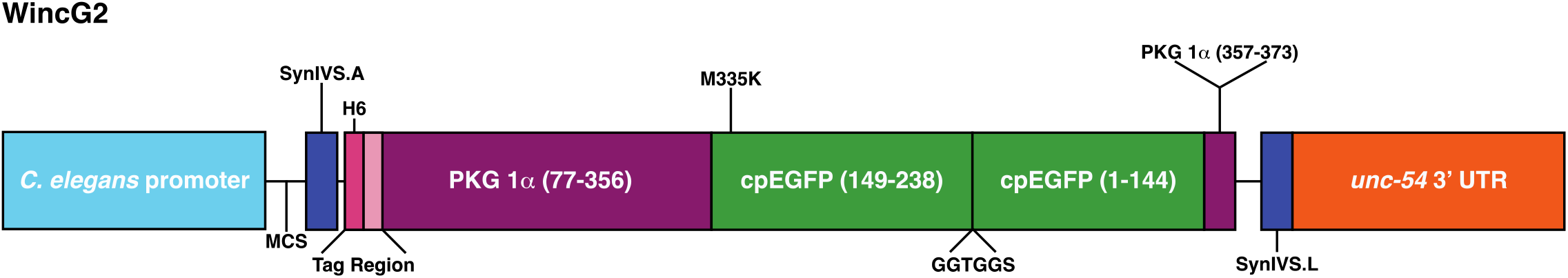
WincG2 fluorescence in the ASER cell body changes linearly in response to 50 mM NaCl step changes and depends on the receptor guanylyl cyclase GCY-22. (A) Average fluorescence response (ΔF/F_0_) of WincG2 in ASER cell body shown as traces responding to either 10 second steps between 50 and 0 mM NaCl or switch control (represented at the bottom of the panel). WincG2 fluorescence in wild-type (N2) animals decreases linearly in response to a 50 to 0 mM NaCl downstep and stops decreasing in response to a 0 to 50 mM NaCl upstep (blue trace, bottom). These responses are not seen in *gcy-22(tm2364)* animals (teal trace, middle) or when wild-type animals are exposed to switch control (pink trace, top). n=17, n=23 and n=11 for wild-type, *gcy-22(tm2364)* and wild-type switch control, respectively. Regression analysis was applied to the data for the first 50 to 0 mM NaCl downstep. R^2^ = 0.9962, R^2^ = 0.0436 and R^2^ = 0.1515 for wild-type, *gcy-22 (tm2364)* and wild-type switch control, respectively. (B) WincG2 fluorescence in the ASER cell body decreases in response to a 50 to 0 mM NaCl downstep and stops decreasing in response to a 0 to 50 mM NaCl upstep in wild-type animals. The slopes for the first 50 to 0 mM NaCl downstep between wild-type and *gcy-22(tm2364)* animals are different (n = 17 (first set, blue; wild-type), n=23 (fifth set, teal; *gcy-22*); permutation test p<0.00001). In wild-type animals, the slopes in response to the first 50 to 0 mM NaCl down step are also different from those of the switch control (n= 17 (first set, blue; 50 mM NaCl downstep), n=11 (third set, pink; switch control); permutation test p<0.00001). The slope values between the first 50 to 0 mM NaCl downstep and 0 to 50 mM NaCl upstep are different in wild-type animals (n=17; first pair, blue; permutation test p<0.00001), as compared to those of the switch control in wild-type animals and *gcy-22* animals, which were not different (n=11; second pair, pink and n=23; third pair, teal, respectively). See Materials and Methods for details of statistical analysis. (C) WincG2 fluorescence in the ASER cell body increases in response to the second 0 to 50mM NaCl upstep in wild-type animals. The slopes for the second 0 to 50 mM NaCl upstep between wild-type and *gcy-22(tm2364)* animals are different (n = 17 (blue; wild-type), n=23 (teal; *gcy-22*); permutation test p<0.05). In wild-type animals, the slopes in response to the second 0 to 50 mM NaCl upstep are also different from those of the switch control (n= 1T (blue; 50 mM NaCl downstep), n=11 (pink; switch control); permutation test p<0.0001).

To test whether changes in WincG2 fluorescence were due to changing NaCl concentrations or to the potential fluctuation in pressure due to the change in flow of the stimulus presentation stream, we examined the sensor’s responses to ten 10-second switches of 50 mM NaCl. WincG2 fluorescence did not change in these animals in response to switching (Fig 3A: pink trace, top; Fig 3B: second pair, pink). Additionally, the slopes of the first downstep between animals recorded in response to changing NaCl concentration and animals recorded in response to simply switching the buffer stream are different (p<0.00001), suggesting that WincG2 fluorescence changes were due to NaCl concentration steps and not to fluctuations in fluid pressure on the exposed nose of the animal (Fig 3B).

WincG2 has a cGMP-binding motif that could also potentially accommodate cAMP, albeit with lower affinity [23,27]. To assess whether WincG2 fluorescence was dependent on cGMP or cAMP, we recorded ASER WincG2 fluorescence in animals lacking the rGC GCY-22. Though other rGCs are expressed in ASER [3,16], loss of GCY-22 produces the most severe behavioral defects in Cl^−^ and NaCl chemotaxis [3]. Consistent with these findings and in contrast to wild-type animals, WincG2 fluorescence did not change in *gcy-22(tm2364)* animals in response to NaCl downsteps or upsteps (Fig 3A: teal trace, ns; Fig 3B: third pair, teal). This finding indicates that changes in WincG2 fluorescence require GCY-22 and likely result from changes in cGMP rather than cAMP.

Wild-type WincG2 fluorescence also increased relative to switch control and *gcy-22* animals in response to the second, third and fourth 0 to 50 mM NaCl upstep (Fig 3C and S2 Fig) and was dependent on both an actual change in NaCl concentration (Fig 3C, compare blue trace with pink trace, p<0.0001) and the rGC GCY-22 (Fig 3C, compare blue trace with teal trace, p<0.05). Together, these results suggest that WincG2 can report rapidly changing increases and decreases in endogenous cGMP levels.

### Animals expressing WincG2 in ASER prefer higher NaCl concentrations relative to animals that do not express the reporter

*C. elegans* requires ASER activity to adjust their preferred NaCl concentration to the concentration at which they were last fed; if ASER is killed, the animal’s movement is less directed in response to a linear NaCl gradient [26]. Plasticity requires NaCl sensation, which in turn requires cGMP signaling; thus it is not surprising that *gcy-22(tm2364)* animals which we showed to not respond to NaCl concentration changes (Fig 3) do not exhibit a preference for the concentration of NaCl at which they were cultivated [16]. To assess whether WincG2 expression affected an animal’s ability to exhibit a preference for its cultivation NaCl concentration, the behavior of WincG2-expressing wild-type animals were compared to their nontransgenic siblings that did not express the WincG2 array and wild-type animals. Animals were cultivated for approximately six hours in the presence of OP50 *E. coli* bacteria on an NGM plate containing 25 mM, 50 mM, or 100 mM NaCl, then placed onto a chemotaxis assay plate containing a NaCl gradient from approximately 40 to 90 mM NaCl (Fig 4A, adapted from [16]). A chemotaxis index (CI) of 1 indicates the animals’ preference for the higher NaCl concentration, and a CI of −1 indicates the animals’ preference for the lower NaCl concentration. Wild-type and nontransgenic siblings behaved as previously described; animals that were cultivated at 25 mM, 50 mM, and 100 mM NaCl had a CI approaching −1, 0 and 0.75, respectively (Fig 4B: first and second set of three bars, respectively) [16]. The WincG2-expressing animals’ NaCl concentration preference at each cultivation NaCl concentration was higher, though not significantly different from wild-type animals (Fig 4B: third set of three bars); however, their preference for a higher NaCl concentration was significantly different from their nontransgenic siblings when they were cultivated at 100mM NaCl (p<0.05). Additionally, the animals’ preference for a higher NaCl concentration seemed different from their nontransgenic siblings when they were cultivated at 50mM, though this was not significant (p=0.15), presumably due to variability of the transgenic animals’ chemotaxis to NaCl when cultivated at 50mM. Other lines injected at the same concentration exhibited NaCl seeking behavior that was significantly different from both wild-type animals and their nontransgenic siblings (S3 Fig). This may indicate that WincG2 expression lowers free cGMP levels and therefore interferes with an aspect of cGMP dynamics in ASER that is required for food to reset the animals’ preference to their cultivation NaCl concentration.

**Fig 4.**
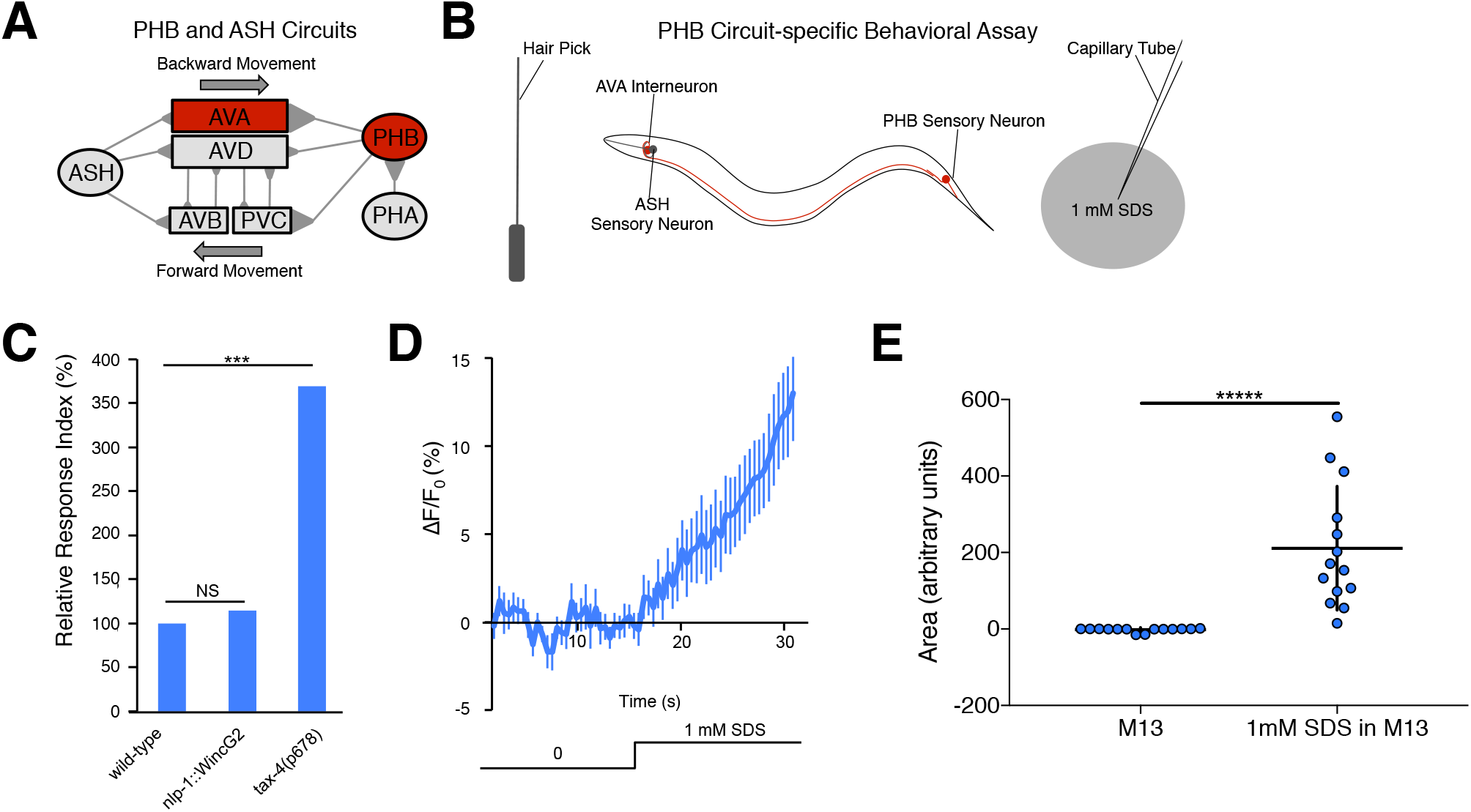
WincG2 expression in ASE does not affect reporter activity but can increase NaCl seeking behavior. (A) Animals were tested for NaCl cultivation preference adapted from Iino [18]. Briefly, animals were placed on cultivation plates with various NaCl concentrations for around six hours before being placed on an assay plate with regions of higher and lower NaCl concentration. (B) Wild-type animals injected with 15 ng/μl *flp-6p*::WincG2 exhibited a preference for higher [NaCl] while maintaining behavioral plasticity; animals from this line were recorded and analyzed for panels C, D, and E. Wild-type animals cultivated at 25mM NaCl, 50mM NaCl, and 100mM NaCl have a chemotaxis index (CI) approaching −1, 0 and 0.75, respectively (first three bars). The line injected with 15 ng/μl *flp-6p*::WincG2 produces two types of progeny: those that do not express the array (nontransgenic siblings) and those that do (transgenic siblings). Nontransgenic siblings behave like wild-type animals (second three bars) (Welch’s t test, ns). Transgenic animals cultivated at each NaCl concentration exhibit a slight but not significant preference for higher NaCl concentration relative to wild-type animals (Welch’s t test, ns). (C) Average fluorescence response ΔF/F_0_) of WincG2 in ASER cell body shown as traces responding to ten 10 second steps between 50 and 0 mM NaCl (represented at the bottom of the panel). WincG2 fluorescence in wild-type (N2) animals decreases linearly in response to a 50 to 0 mM NaCl downstep and stops decreasing in response to a 0 to 50 mM NaCl upstep; this is seen in animals injected with either 7.5 ng/μl (top trace; blue) or 15 ng/μl (bottom trace; red) *flp-6p*::WincG2. The data for the strain injected with 7.5 ng/μl *flp-6p*::WincG2 are the same data that was used in Fig 4A as wild-type (N2). Regression analysis was applied to the data for the first 50 to 0 mM NaCl downstep. R^2^ = 0.9962 and R^2^ = 0.9963 for wild type animals injected with 7.5 ng/μl and 15 ng/μl *flp-6p*::WincG2, respectively. (D) The slope values of the first 50 to 0 mM NaCl downstep and 0 to 50 mM NaCl upstep are different in wild-type animals injected with either 7.5 ng/μl (n=17; first pair, blue; permutation test p<0.00001) or 15 ng/μl (n=18; second pair, red; permutation test p<0.00001) *flp-6p*::WincG2. The data for the strain injected with 7.5 ng/μl *flp-6p*::WincG2 are the same as the data used in Fig 4B as wild-type, first pair, blue. The slopes for the first 50 to 0 mM NaCl downstep in wild-type animals injected at 7.5 ng/μl and 15 ng/μl *tlp-6p*::WincG2 are not different (n = 17 (blue; 7.5 ng/μl, n=18 (red; 15 ng/μl; permutation test ns).

WincG2 fluorescence was recorded for the line that exhibited behavior that was not significantly different from wild-type animals. Importantly, in these animals, WincG2 fluorescence decreased linearly in response to a 50 to 0 mM NaCl downstep and stopped decreasing in response to a 0 to 50 mM NaCl upstep, with the slopes between the first downstep and upstep being different (Fig 4C: red trace, bottom; Fig 4D: second pair, red; p<0.00001). This finding is similar to that observed with WincG2 injected at a lower concentration (Fig 4D: first pair, blue; note that these data points are reproduced from Fig 3A: blue trace, bottom and Fig 3B: first pair, blue, respectively). The slopes of the first downstep of wild-type animals injected at a lower versus higher concentration are not different, indicating that the concentrations injected did not influence recordings (Fig 4D: compare first blue trace with third red trace; p ns). Thus, though WincG2 reliably reports the stimulus-induced changes in the gustatory sensory neuron ASER, its expression may subtly alter behavior and this must be controlled for by comparing behavior in transgenic lines to the non-transgenic siblings.

### WincG2 fluorescence increases in PHB chemosensory tail neurons in response to SDS

To examine the ability of WincG2 to report cGMP changes in a neuron with a different modality, we expressed WincG2 in the nociceptive PHB neurons, which had been predicted to use cGMP as a second messenger. The PHBs are a pair of bilaterally symmetric sensory neurons located in the lumbar ganglia that extend ciliated dendrites into the phasmid structures within the tail of *C. elegans*. PHB neurons are chemosensory cells that are required for the avoidance of noxious chemicals such as SDS, dodecanoic acid and other cues (Fig 5A) [19,20,32,33]. TAX-4 is required for PHB-mediated SDS avoidance (Fig 5C), and calcium imaging has shown that PHB responds to SDS [20]. This suggests that PHB may exhibit changes in cGMP levels in response to SDS that could be monitored by recording changes in WincG2 fluorescence.

**Fig 5.**
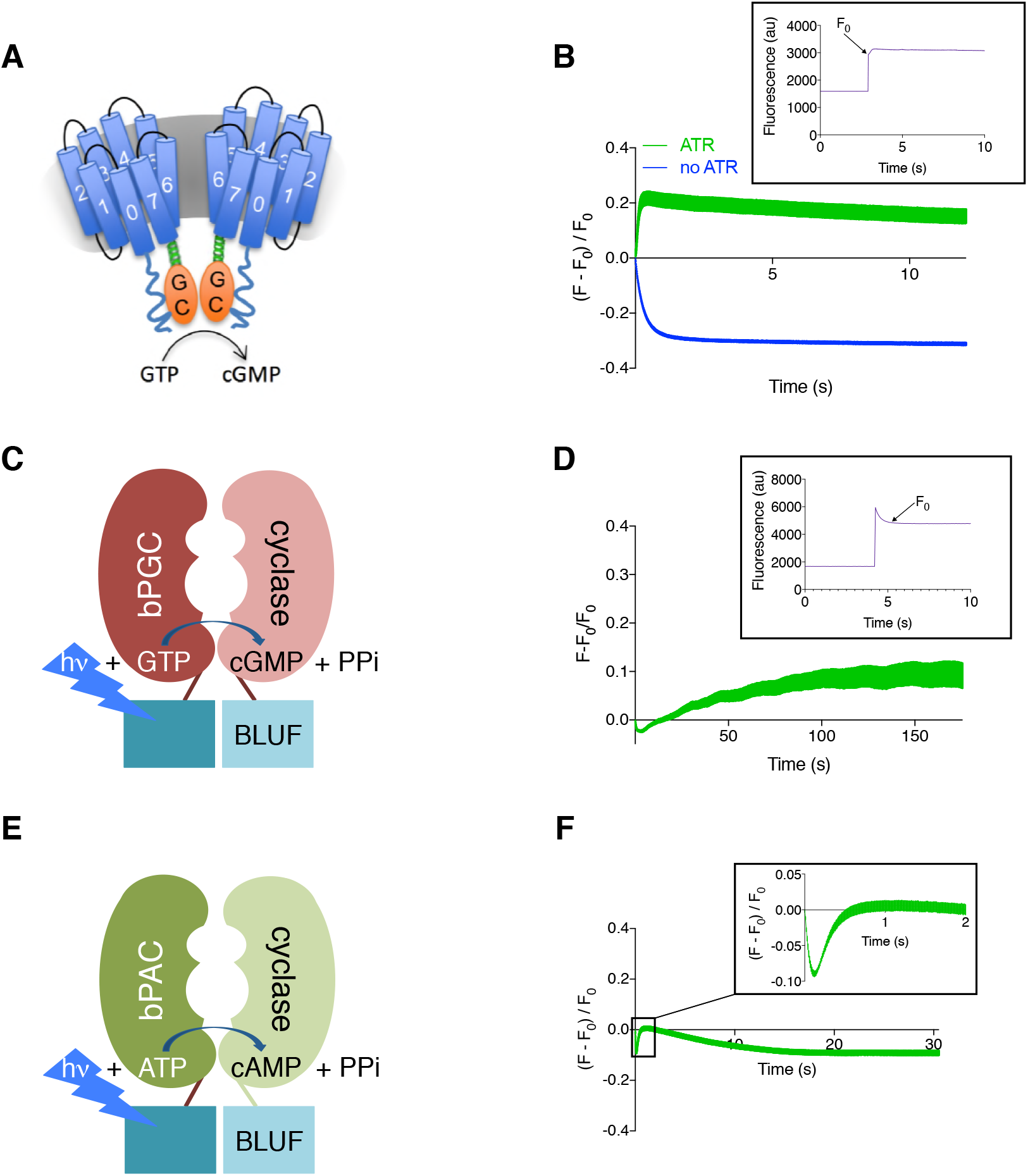
WincG2 allows visualization of cGMP production in PHB chemosensory neurons in response to the repellent SDS. (A) Diagram of the PHB circuit [19,32,33]. The primary postsynaptic partners of PHB neurons are the AVA backward command interneurons and the PVC forward command interneurons. (B) Diagram of the sodium dodecyl sulfate (SDS) behavioral assay, which tests the PHB circuit response to 1mM SDS [19,32,33]. Animals are induced to move backwards with a nose touch from a hair pick. A drop of 1 mM SDS is placed behind them on a dry NGM plate, so that the drop quickly absorbs into the media, preventing wicking along the worm. The time that the animal backs into the drop before stopping is measured. (C) WincG2 does not affect the response of animals to SDS. For reference, loss of *tax-4* cyclic nucleotide-gated channel causes a severe defect in the ability to sense SDS [32]. (D) When WincG2 is expressed in PHB neurons, AF/F_0_ increases steadily after introduction of 1 mM SDS. n=16 (E) The area under the curve for traces before and after SDS presentation is different (permutation test, p<−0.00001).

To test whether WincG2 affects the function of the PHB circuit, SDS response assays (Fig 5B) were performed on wild-type animals and animals expressing WincG2 in PHB neurons. On average, wild-type animals halt movement into a drop of 1 mM SDS in less than a second. If PHB function is impaired, as in *tax-4* mutants, animals continue moving into a drop of SDS as if it were a control buffer (M13). This increases the relative response index to approximately 300% (Fig 5C). We found that behavioral responses to SDS were unaffected by WincG2 expression, indicating that WincG2 does not affect PHB function (Fig 5C).

To determine if cGMP changes in PHB neurons could be detected using WincG2, the sensor’s fluorescence in the cell body was measured in animals that were first exposed to control buffer (M13), then to 1 mM SDS in M13 buffer. WincG2 fluorescence remained largely steady in the absence of 1 mM SDS, but began to increase linearly (R^2^=0.9633) upon exposure to 1 mM SDS (Fig 5D). The area of the curve for the traces before and after SDS presentation is different (permutation test, p<0.00001), which suggests that cGMP increases in response to SDS and that WincG2 responds acutely to endogenously produced cGMP that is induced by an external stimulus (Fig 5E).

## Discussion

### WincG2 is a sensor for cGMP dynamics in *C. elegans*

The cGMP sensor WincG2 was successfully used to monitor the dynamics of this second messenger in a number of cells in *C. elegans*. First, WincG2 was used to monitor the kinetics of cGMP production in body wall muscles that lack most endogenous GCs. The rate of increase in WincG2 fluorescence corresponded with the rate of cGMP produced by coexpressed blue light-activated GCs. WincG2 fluorescence increased within less than a second of activation of BeCyclOp, which produces 17 cGMP molecules per second. In contrast, WincG2 fluorescence increased in the order of minutes upon activation of bPGC, which produces cGMP at a 50-fold lower rate relative to BeCyclOp [24]. WincG2 fluorescence slightly increased in the presence of cAMP in *C. elegans* in response to activation of bPAC. Thus, care must be taken to control for fluctuations in cAMP by imaging in backgrounds that lack cGMP production. The high rate of cAMP production due to bPAC, however, is likely exceeding any intrinsic cAMP production by several fold, thus side effects from intrinsic cAMP fluctuation may affect cGMP imaging only to a minor extent.

### WincG2 reveals cGMP dynamics in sensory neurons that use cGMP as a second messenger for sensory stimuli

We expressed WincG2 in sensory neurons that use cGMP as a second messenger and found that the sensor responds robustly to changing stimulus presentation. In the gustatory neuron ASER, changes in WincG2 fluorescence in response to repeated NaCl concentration steps suggest that the sensor can respond reliably to changing cGMP levels, providing, for the first time, visual evidence that cGMP levels in ASER are modulated by NaCl concentration changes. Importantly, we demonstrated that WincG2 fluorescence changes require the rGC GCY-22. This has two implications: 1) the sensor responds to cGMP in a physiologically relevant setting and 2) GCY-22 seems to be the primary rGC that produces cGMP for the NaCl response. Interestingly, the changes we observe in WincG2 fluorescence in ASER suggest that cGMP levels may be inversely correlated with calcium in response to a NaCl concentration change. Previously reported genetic and calcium imaging evidence suggested that cGMP levels would directly correlate with calcium levels; cGMP is hypothesized to bind and open the TAX-2/TAX-4 cyclic nucleotide-gated cation channel to allow for calcium influx [25]. A recent cryo-EM study showed that cGMP gates the homomeric TAX-4 channel in the open state [34]. Additionally, physiological investigations of the heterologously expressed heteromeric TAX-2/TAX-4 cation channel indicated that it opens in response to cGMP binding [35,36]. The unexpected result that WincG2 fluorescence decreases in response to NaCl downsteps could be due to the tight regulation of cGMP levels by phosphodiesterases. Resolution of this apparent paradox, however, awaits further investigation.

In the nociceptive phasmid neuron PHB, genetic evidence suggesting that it uses cGMP to signal the presence of SDS was corroborated by changes in WincG2 fluorescence. This is the first visual evidence for a cGMP-based signal in PHB, showing that it increases in response to an environmental cue. Importantly, expression of WincG2 did not perturb the function of the phasmid neurons, as SDS repulsion was as robust in the WincG2-expressing line as in wild-type animals. This is in contrast to WincG2 expression in ASER, which caused a slight preference for higher NaCl concentrations but minimally affected the plasticity of the NaCl concentration cultivation preference.

### Prospects for optimizing WincG2 and extending its use

Like WincG2, the first generation of calcium sensors affected the behavior of neurons in which they were expressed: the FRET-based YC2.12 acted as a calcium sponge, a function that was exploited by Ferkey et al. to study nociceptive signaling [37], and the GFP-based GCaMP2.2 blocked olfactory plasticity (personal communication, Brueggemann and L’Etoile). Mutations that increased the quantal yield of GCaMP allowed the reporters to be expressed at lower levels that did not interfere with cellular function, yet were bright enough for imaging. Indeed, if WincG2 was enhanced to mimic the properties of GCaMP6s, which contains (among other mutations) a K78H mutation in the cpEGFP domain that improved sensitivity relative to GCaMP3, the fluorescence might be bright enough to allow for lower expression of this reporter and thus reduce the possibility of it interfering with cellular functions [38]. Until such optimizations are made, it will be necessary to select for lines that express WincG2 at the lowest levels that allow for imaging thereby minimizing the potential for behavioral effects. Addition of a subcellular localization signal may also mitigate off-target effects.

We think WincG2 could be acting as a cGMP sponge due to its effects on NaCl-seeking behavior when expressed in ASE (Fig 5B). These behavioral results suggest that WincG2 could be altering free cGMP levels in ASER, which could lead to tuning the NaCl concentration cultivation preference to be higher relative to nontransgenic siblings and wild-type animals. If WincG2 can be shown to act as a cGMP sponge, this could also be exploited to specifically and locally perturb cGMP levels. For example, if one could localize a non-fluorescent form of WincG2 at the cilia, this may perturb function in a different way from when it is localized to the cell body. This would reveal specific functions for cGMP signals at the sensory dendrites that are different from those in the cell body.

Additionally, the subcellular landscape of cGMP can also be probed using WincG2. For instance, adding a small tag that localizes WincG2 to specific regions of the cell along with a red protein for ratiometric imaging may reveal important aspects of the subcellular landscape of cGMP.

Though FRET-based cGMP sensors have been useful for uncovering cGMP dynamics in biological processes in an intact animal [8,22], the single fluorophore WincG2 provides the possibility of using additional fluorophores of different wavelengths to mark subcellular regions. This advantage of WincG2 will provide a simple and powerful tool with which to visualize changes in cGMP concentration across the subcellular landscape. Additionally, the ability to simultaneously visualize cGMP and calcium by using WincG2 with a red calcium sensor will allow us to investigate the dynamics of these second messengers at any marked subcellular location [39]. Our results demonstrate that WincG2 can be used to rapidly and specifically measure cGMP dynamics in the intact, behaving organism.

## Materials and Methods

### Molecular Biology

FlincG3 was codon optimized by Genscript for use in *C. elegans*. The *C. elegans* codon-optimized FlincG3, WincG2, was inserted into a worm specific expression (Fire) vector, pPD95.75, which facilitates transcription and translation of the sensor by providing worm specific introns [40].

The *gcy-5* promoter (2003 bp upstream of start site) was inserted using XbaI and EagI into pPD95.75 to make *gcy-5p*::WincG2. *myo-3p*::CyclOp::SL2::mCherry was made as described [24]. The plasmid *gcy-5p*::WincG2 was digested with XbaI and PciI and the *gcy-5* promoter fragment was replaced with the *myo-3* promoter fragment to yield the *myo-3p*::WincG2 plasmid construct.

pCFJ104 [*myo-3p*::mCherry::unc-54] was a gift from Erik Jorgensen (Addgene plasmid # 19328) [41]. To generate *myo-3p*::bPAC::SL2::mCherry, a 1193 bp long bPAC fragment was amplified using primers 5’-TCCATCTAGAGGATCCTTCCGCATCTCTTGTTCAAGGG-3’ and 5’-CAGCGGTACCGTCGACTTACTTGTCGTTTTCCAGGGTCTG-3’. This was ligated into *myo-3p*::SL2::mCherry backbone, obtained by digestion with BamHI and SaII, using ‘in-fusion cloning’ (Clontech). To generate *myo-3p*::bPGC::SL2::mCherry, a 3486 bp long *myo-3p::bPGC* fragment was amplified using primers 5’-ATTACGCCAAGCTTGCGGCTATAATAAGTTCTTGAA-3’ and 5’-TACCGTCGACGCTAGTTACTTGTCGTTTTCCAGGG-3’. This was ligated into a SL2::mCherry backbone, obtained by digestion with NheI and SphI, using ‘in-fusion cloning’ (Clontech).

To generate the *nlp-1p*::WincG2 construct, WincG2 from *gcy-5p*::WincG2 was inserted into the vector *nlp-1p*::pSMΔ [42] using the NEBuilder High-Fidelity DNA Assembly Cloning Kit (NEB) and the following primers: MVP598 WincG FWD (5’-ggattggccaaaggacATGGCACACCACCACCAC-3’), MPV599 WincG REV (5’-ggtcctcctgaaaatgttcTTATCGTCCGAATCCTCCG-3’), MVP597 *pSM*Δ*::nlp-1p* FWD (5’-GAACATTTTCAGGAGGACCC-3’), and MVP596 *nlp-1p::pSM*Δ REV (5’-GTCCTTTGGCCAATCCCG-3’). The construct was then sequence-verified.

To generate the *flp-6p*::WincG2 construct, the *flp-6* promoter fragment was amplified from *flp-6p*::GCaMP6, a gift from the Lockery lab, using the primers 5’-ATTACGCCAAGCTTGCATGGCAGCGCTTGACTTCTGATG-3’ and 5’-CCGGGGATCCTCTAGTGCAGGCATGCAAGCTTGTC-3’. The plasmid *nlp-1p*::WincG2 was digested with XbaI and SphI-HF and the *nlp-1* promoter fragment was replaced with the *flp-6* promoter fragment to yield the *flp-6p*::WincG2 plasmid construct using ‘in-fusion’ cloning (Clontech). The plasmid str-2::jRCaMP1b, a gift from the Bargmann lab, was digested with HindIII-HF and XmaI and str-2 was replaced with the *flp-6* promoter fragment to yield the *flp-6p*::jRCaMP1b plasmid construct using ‘in-fusion’ cloning (Clontech). The plasmid *str-2p*::jRGECO1a, a gift from the Bargmann lab, was digested with HindIII-HF and AscI and the *str-2* promoter fragment was replaced with the *flp-6* promoter fragment to yield the *flp-6p*::jRGECO1a plasmid construct using ‘in-fusion’ cloning (Clontech).

pJN55 [*myo-3p*::tax-2::GFP] and pJN58 [*myo-3p*::tax-4::GFP] plasmid construction have been previously reported [24]. pWSC13 [*myo-3p*::bPGC::eYFP]: pPD96.52 was cut with NheI and BamHI to yield a 6010 bp fragment, and bPGC synthetic construct, a gift from Peter Hegemann, was cut with BamHI and NheI to yield a 2106 bp fragment that was then inserted into cut pPD96.52. pJN59 [*myo-*3p::bPGC(K265D)::eYFP]: K265D was introduced into the parent plasmid pWSC13 (*myo-3p*::bPGC::eYFP) using the Q5 site-directed mutagenesis kit (New England Biolabs Inc.) and primers oJN146 (5’-CCTGAAGATGGACCACGGCCTGC-3’) and oJN147 (5’-CTGCTGCCCATGTTGCCC-3’).

### Transgenic Strains

Transgenic *C. elegans* were obtained by microinjection of DNA into the gonads of nematodes by standard procedures [43]. *ZX1921 (zxEx895[myo-3p::CyclOp::SL2::mCherry, myo-3p*::*WincG2]):* 15ng/μl *myo-3p*::CyclOp::SL2::mCherry and 15ng/μl *myo-3p*::WincG2 were microinjected into N2 background worms. *ZX1757 (zxEx893[myo-3p::mCherry, myo-3p*::*WincG2]):* 5 ng/μl *myo-3p*::mCherry and 20 ng/μl *myo-3p*::WincG2 were microinjected into N2 background worms. *ZX1922 (zxEx896[myo-3p::bPAC::SL2::mCherry, pmyo-3*::*WincG2]):* 15ng/μl *myo-3p*::bPAC::SL2::mCherry and 15ng/μl *myo-3p*::WincG2 were microinjected into N2 background worms. *ZX1756 (zxEx892[myo-3p::bPGC::SL2::mCherry, myo-3p*::*WincG2]):* 15ng/μl *myo-3p*::bPGC::SL2::mCherry and 15ng/μl *myo-3p*::WincG2 were microinjected into N2 background worms. *JZ1997 (ASE WincG2 in N2 injected with 15ng/μl WincG2):* 15ng/μl *flp-6p*::WincG2, 30 ng/μl *flp-6p*::jRCaMP1b and 20ng/μl ofm-1::GFP were microinjected into N2 background worms. *JZ2089 (ASE WincG2 in N2 injected with 7.5 ng/μl WincG2):* 7.5ng/μl *flp-6p*::WincG2, 60 ng/μl *flp-6p*::jRGECO1a and 20ng/μl ofm-1::GFP were microinjected into N2 background worms. *JZ2118 (ASE WincG2 in gcy-22):* JZ2089 animals were crossed with OH4839 *(gcy-22(tm2364))* animals to generate transgenic animals homozygous for *gcy-22(tm2364). PHB WincG2 in N2:* MKV937 *(iyEx222* [15ng/μL *nlp-1p::WincG2* and 45ng/μl *odr-1p::RFP* in N2 background worms]). *ZX1738 (zxEx886[myo-3p::bPGC::YFP, myo-3p::tax-2::GFP, myo-3p::tax-4::GFP]):* 15ng/μl *myo-3p*.:bPGC::YFP, 5 ng/μl *myo-3p*:tax-2::GFP and 5 ng/μl *myo-3p*:tax-4::GFP were injected into *lite-1(ce314)* background worms. *ZX1739 (zxEx887[myo-3p::bPGC(K265D):: YFP, myo-3p::tax-2::GFP, pmyo-3::tax-4::GFP]):* 15ng/μl *myo-3p*:bPGC(K265D)::YFP, 5 ng/μl *myo-3p*.:tax-2::GFP and 5 ng/μl *myo-3p*::tax-4::GFP were injected into *lite-1(ce314)* background worms.

### Imaging WincG2 coexpressed with BeCyclOp, bPGC or bPAC

Transgenic strains were kept in the dark on standard NGM plates (5.5 cm diameter; 8ml NGM) with OP50–1 bacteria with or without ATR at 20°C. Plates containing ATR were prepared by spreading 200 μl of OP50–1 culture containing 100 mM of ATR (diluted in ethanol). L4 animals were put on ATR plates overnight and young adults were used for imaging the following day.

For cGMP/cAMP imaging, animals were immobilized on 10% M9 agar pads with polystyrene beads (Polysciences, USA). The fluorescence measurements were performed with a 25x oil objective (Zeiss 25x LCI-Plan / 0.8 Imm Corr DIC) on the inverted microscope Axio Observer Z.1 equipped with two high-power light emitting diodes (LEDs; 470 and 590 nm wavelength, KSL 70, Rapp Optoelektronik, Germany) coupled via a beam splitter and a double band pass excitation filter permitting wavelengths of 479/21 nm and 585/19 (F74–423 479/585 HC Dualband Filter AHF Analysentechnik) to obtain simultaneous dual-wavelength illumination. DIC microscopy using white light filtered with a red optical filter was used to focus on the body wall muscle cells prior to video acquisition. The 470nm and 590 nm excitation was switched on simultaneously after the start of video acquisition. For bPGC experiments, yellow light was used to focus the cells, and thereafter the blue illumination was turned on. Fluorescence was acquired by an ORCA-Flash 4.0 sCMOS camera (Hamamatsu) through a Dual-View beam splitter (DV2, Photometrics) with a 540/25 nm emission filter used for WincG2 green channel and a 647/57 emission filter for mCherry red channel. Videos were acquired using μManager [44], and frames were taken at 100Hz (corresponding to exposure times of 10 ms) and 20Hz (for bPGC) with 4×4 spatial binning. The optical power was 3.3 mW/mm^2^ at 470 nm and 2.6 mW/mm^2^ at 590 nm.

Image analysis was performed in ImageJ (NIH). Regions of interest (ROIs) were drawn around single body wall muscle cells that did not show major movement and a region outside the animals was chosen as background ROI. The mean fluorescence intensity of the ROIs for both channels was analyzed with ImageJ. Background subtracted values were used to calculate the change in fluorescence intensity for each time point: (F-F_0_)/F_0_ where F represents the intensity at this time point and F_0_ represents the peak intensity at the onset of light stimulation. For bPGC experiments, F_0_ represents the intensity 5s after the onset of light stimulation.

### Imaging WincG2 in ASER and PHB

WincG2 imaging was performed essentially as described for calcium imaging [17]. Briefly, for imaging ASER, day 1 adults grown at 20°C were transferred from NGM plates containing OP50 to a 35 mm × 10 mm petri dish containing chemotaxis buffer with 50 mM NaCl (25 mM K_3_PO_4_, pH 6.0, 1 mM CaCl_2_, 1 mM MgSO_4_, 50 mM NaCl, adjusted to 355 ± 2 mOsm with sorbitol). The worms were then placed in a microfluidic device that can expose the animal to stimulus [45]. A Zeiss 40x air objective on an inverted microscope (Zeiss Axiovert 200) was used for imaging, and images were taken at rate of 1Hz with a blue light exposure time of 30 ms using an ORCA-Flash 2.8 camera (Hamamatsu) for a total of 103 frames. Recordings were taken within eight minutes of the animal’s exposure to chemotaxis buffer with 50 mM NaCl, and the animals were subjected to either ten second steps between chemotaxis buffer with 50 and 0 mM NaCl (each containing 1 mM levamisole) or switches between chemotaxis buffer with 50 mM NaCl (each containing 1 mM levamisole (Sigma-Aldrich), and one containing fluorescein). Images were obtained using μManager (Version 1.4.22). Fluorescence intensity was measured using ImageJ. To calculate the fluorescence intensity at a given time point (F), the fluorescence intensity from the region of interest (ROI) encompassing the neuron was subtracted from the background ROI. The fluorescence intensity F of the first three frames was averaged to calculate F_0_. We used ΔF/F_0_ (F-F_0_/F_0_) to calculate the change in fluorescence intensity at a given time point.

To image WincG2 in PHB neurons, animals were picked from NGM plates containing OP50 onto a petri dish containing M13 control buffer, then placed tail-first into the microfluidic device. They were exposed to control buffer for 15 seconds, and then to 1 mM SDS in M13 for 15 seconds. They were imaged at a rate of 2 Hz. To calculate the fluorescence intensity of PHB at each frame, Image J was used to measure the total intensity of the cell body. Background fluorescence was calculated by using ImageJ to measure the minimum pixel value in the area surrounding the cell body, and this pixel value was multiplied by the area of the cell body to get the total background. The total background was subtracted from the total intensity of the cell body to calculate the fluorescence intensity. The PHB WincG2 fluorescence intensities were adjusted for photobleaching using the following method. The decrease in fluorescence during the first 29 frames when the animal was exposed to control buffer was presumed to be due to photobleaching. Therefore, the average difference between the values for the nth frame and the n+1th frame (up to the 29^th^ frame) were calculated, and this average photobleaching value was then added back to each value in the series. F_0_ was the average of the response to buffer adjusted for photobleaching over the first 15 seconds, rather than only the first 3 data points.

### NaCl cultivation assay

The NaCl cultivation assay was essentially performed as described [16]. Briefly, day 1 animals grown at 20°C were transferred from OP50-containing NGM plates containing 50 mM NaCl to OP50-containing NGM plates containing 25 mM, 50 mM, or 100 mM NaCl for approximately six hours before being placed on a chemotaxis assay plate containing regions of higher and lower NaCl for 45 minutes. Afterward, worms were stored in 4°C for at least 16 hours before calculating the chemotaxis index. Chemotaxis index = [# animals at higher NaCl region − # animals at lower NaCl region]/ [# animals at higher NaCl region + # animals at lower NaCl region + # animals outside origin].

### Behavioral assay response to the repellent SDS

SDS dry drop behavioral assays were conducted by using a hair pick to touch each worm on the nose to stimulate backward movement into a dry drop of 1 mM SDS in M13 buffer [32,33]. A dry drop is obtained by incubating an NGM plate overnight at 37°C so that the SDS drop dries quickly into the plate, preventing wicking along the animal that might activate neurons in the head. An animal’s response time was defined as the amount of time it backed into the dry drop before terminating backward movement. The average response time to the dry drop of dilute SDS in M13 buffer was compared to the average response time to a drop of control M13 buffer. A response index was calculated by dividing the average response time to SDS by the average response time to M13 buffer. *nlp-1p*::WincG2 and *tax-4* mutants were each compared to wild-type animals assayed on the same day, and the wild-type response index is normalized to 100%. At least 80 animals of each genotype were tested: 40 for a response to M13, and 40 for a response to SDS.

### Muscle Contraction Assay

The muscle contraction assay was essentially performed as described [24]. L4 animals were exposed to 0.9 mW/mm^2^ blue light (450–490 nm) for approximately 20 seconds, and relative body length was measured with a custom LabView script.

### Statistical Analysis for ASER and PHB WincG2

To compare the sum of slope values of the first 50 to 0 mM NaCl downstep with the first 0 to 50 mM NaCl upstep, as well as the sum of slope values of the first 50 to 0 mM NaCl downstep and the second, third, and fourth 0 to 50 mM NaCl upstep, an approximate p-value ranked permutation test was performed on the data. To compare the sum of slope values of the first 50 to 0 mM NaCl downstep with the first 0 to 50 mM NaCl upstep, the sum of slopes for randomly rearranged data is calculated and ranked relative to the sum of slopes for the first 50 to 0 mM NaCl downstep. For example, to compare the sum of slope values of the first 50 to 0 mM NaCl downstep with the first 0 to 50 mM NaCl upstep for wild-type animals, the 34 slope values of the first downstep and upstep were combined in a list then randomly rearranged, and the sum of the first 17 slope values of that randomly rearranged list was calculated and ranked relative to the sum of 17 slope values for the first 50 to 0 mM NaCl downstep. To compare the sum of slope values of the first 50 to 0 mM NaCl downstep between conditions, the sum of slopes for the randomly rearranged data is calculated and ranked relative to the sum of slopes for wild-type. For example, for comparing the sum of slope values of the first 50 to 0 mM NaCl downstep between wild-type and *gcy-22* animals, the 40 (n=17 for wild-type and n=23 for *gcy-22*) slope values were combined in a list then randomly rearranged, and the sum of the first 17 slope values of that randomly rearranged list was calculated and ranked relative to the sum of 17 slope values for wild-type. This type of statistical test was also performed for comparing the second, third, and fourth 0 to 50 mM NaCl upstep between conditions. The sum of slopes from at least 500,000 randomly selected permutations were calculated and ranked using a custom Python script. For comparing the sum of slope values of the first 50 to 0 mM NaCl downstep with the first 0 to 50 mM NaCl upstep for wild-type switch control; for comparing the sum of slope values for the first 50 to 0 mM NaCl downstep between wild-type and wild-type switch control; and for comparing the area under the curve for the first 30 frames of PHB WincG2 (corresponding to M13 buffer) with the next 30 frames of PHB WincG2 (corresponding to 1 mM SDS), an exact p-value ranked permutation test was performed. For this test, the sum of slopes for all possible permutations of the data was calculated and ranked using a custom Python script.

## Author contributions

KA, WSC, MF, SW and MWS prepared reagents. JL, JN, BB-R, MWS, RS, AT and AV performed experiments. SW guided the project and performed experiments. OSH initiated the line of experimentation. AG, NDL and MV conceived the experiments and guided the project. DMF and CB helped guide the project. AG, NDL, JN, MV and SRW wrote the manuscript.

## Acknowledgements

The authors would like to thank Martina Bremer and Saul Kato for statistical consults, as well as Cornelia Bargmann, Peter Hegemann, Erik Jorgensen and Shawn Lockery for reagents. The authors would also like to thank Sarah Nordquist and Piali Sengupta for guidance on writing the manuscript. NDL and MKV were supported by NIDCD R01 DC005991 and NINDS R01 NS087544–01; NDL and DMF by NIDCD R01 DC015758; AG by DFG (Deutsche Forschungsgemeinschaft) grants GO1011/4-2, GO1011 /13-1 and EXC115; SW by Diversity Supplement NS87544 and 5T32DE007306-21.

## Supporting Information Captions

**S1 Fig. Coexpression of bPGC with the cGMP-gated cation channel TAX-2/TAX-4 in body wall muscle cells results in muscle contraction within two seconds upon exposure to blue light (n=10).** bPGC(K265D) contains a point mutation that abolishes guanylyl cyclase activity. Animals coexpressing bPGC(K265D) with TAX-2/TAX-4 did not contract to the same extent (n=12).

**S2 Fig. WincG2 fluorescence in the ASER cell body increases in response to the third and fourth 0 to 50mM NaCl upstep in wild-type animals.** The difference in slopes for the third and fourth 0 to 50 mM NaCl upstep between wild-type and *gcy-22(tm2364)* animals is significant (n = 17 (blue; wild-type), n=23 (teal; *gcy-22*); permutation test p<0.05 for third upstep and p<0.01 for fourth upstep). In wild-type animals, the difference between the slopes in response to the third and fourth 0 to 50 mM NaCl upstep is also significant from those of the switch control (n= 17 (blue; wild-type), n=11 (pink; switch control); permutation test p<0.01).

**S3 Fig. Other lines generated from injecting 15 ng/μl *flp-6p*::WincG2 exhibit a preference for higher [NaCl] while maintaining behavioral plasticity.** This preference is significantly different from wild-type animals and nontransgenic siblings.

